# Closed-loop optical optimization enables patterned retinal stimulation *in vivo* at cellular scales

**DOI:** 10.64898/2026.08.07.742359

**Authors:** Justin Chen, Fengyuanshan Xu, Paul J. Jablonski, Roman Kuranov, Xiaorong Liu, Yang Hu, Cheng Sun, Hao F. Zhang

**Affiliations:** Department of Biomedical Engineering, Northwestern University, Evanston, IL, USA; Department of Mechanical Engineering, Northwestern University, Evanston, IL, USA; Department of Ophthalmology, Stanford University, Stanford, CA, USA; Department of Biology, University of Virginia, Charlottesville, VA, USA

**Keywords:** Neurostimulation, photoreceptors, fundus camera, digital micromirror device, cone cells, vision science

## Abstract

Visual neuroscience requires precise spatiotemporal projection of optical stimulation onto the retina, especially in experimental mouse models. However, *in vivo* patterned stimulation in mice is profoundly hindered by the extreme optical power and severe anatomical aberrations of the eye. Consequently, visual stimulation relies mainly on unverifiable, open-loop approximations that often lack spatial precision. Here, we introduce a closed-loop, spatially modulated stimulation platform that overcomes these barriers. By integrating a digital micromirror device (DMD) with electronically tunable lenses (ETLs) and a real-time, fundus camera-guided focus optimization module, we directly verify the location of patterned stimuli on the retina while dynamically correcting for chromatic and geometric defocus. This platform delivers quantitatively verified static and dynamic patterned stimuli to the living retina with lateral resolutions as fine as 6.7 µm. Guided by ray-tracing optical analysis, our work establishes a technological foundation that enables highly reproducible, cellular-scale interrogations of the visual pathway.

## 1 Introduction

Patterned stimulation is a foundational technique for probing visually evoked neuronal activity and function [1, 2]. Unlike full-field flashes, which indiscriminately activate large neuronal populations, patterned stimuli enable the interrogation of receptive-field structure, local contrast sensitivity, and fine spatial processing across retinal output pathways [3]. Previously, researchers used diverse spatiotemporal patterns to examine specific computations, including contrast-reversing checkerboards to assess linearity and drifting bars to quantify direction selectivity of retinal ganglion cells (RGCs) [4]. However, most studies relied on *ex vivo* flat-mount retinas, which lack the physiological and behavioral context of the intact visual system. *In vivo* approaches address this limitation by linking retinal activity to visually guided behaviors, such as the optomotor response, and by enabling electrophysiological assays, including pattern electroretinography, to assess visual pathway integrity [5, 6]. These capabilities are critical for investigating early indications of neurodegenerative diseases, such as glaucoma, during which the functional identities of retinal neuronal subtypes may evolve [7].

Extending patterned stimulation to the intact eye, however, faces significant optical challenges. Image formation on the retina is influenced by the native ocular optics, where variability in refractive power across subjects can degrade image quality [8]. In mice, these effects are exacerbated by the extreme optical power and retinal curvature [9, 10], which introduce both global and field-dependent defocus [8, 11]. As a result, even minor misalignments can attenuate high spatial-frequency components and distort the intended pattern [12].

Despite these constraints, existing *in vivo* stimulation approaches lack mechanisms to verify that the stimulus is correctly projected onto the retina. Many studies place external monitors in front of the subject [13–15], and others employ more intricate systems that use spatial light modulators (SLMs) to project infinity-corrected patterns through the pupil [16]. However, all of these works rely on intact optics to relay the image to the retinal plane, forming an open loop that assumes accurate conjugation between the desired patterns and their images on the retina. Consequently, researchers were unable to verify whether the stimuli were fully passing through the pupil, determine their precise retinal location, or confirm that they were in focus. If this assumption fails, edge-features may become blurred, leading to unintended activation of adjacent neural populations [17].

This technical gap highlights the need for a verifiable patterned stimulation system that adapts to the optical properties of each eye, rather than relying on a predefined, “one-system-fits-all” approach. To this end, we developed a fundus camera-guided platform equipped with a closed-loop feedback mechanism to verify the patterned stimuli on the retina while optimizing stimulus focus. Spatial patterning was achieved using a digital micromirror device (DMD), an SLM comprising a dense array of independently switchable mirrors [18]. Modern DMDs provide sufficient spatial sampling for cellular-scale optical stimulation while supporting refresh rates [19] that exceed biologically relevant timescales, such as the critical flicker fusion frequency (∼ 30 Hz) [20, 21] and RGC firing rates (up to ∼100 Hz) [22]. Integrating a fundus camera enables real-time observation of stimulus patterns, while a focus metric continuously optimizes image quality, ensuring consistent and reproducible pattern delivery across subjects and experimental sessions. This feedback is implemented through a pair of electronically tunable lenses (ETLs), which dynamically adjust the focal planes of both the imaging and stimulation pathways [23].

## 2 Results

### 2.1 Experimental system

Our experimental system consists of three coordinated paths: fundus illumination, patterned stimulation, and fundus imaging, as shown in **Fig. 1(a)**, with a SolidWorks model provided in **Fig. S1**. Additional details of the optical components, including part numbers, are provided in Materials and Methods.

**Figure 1.**
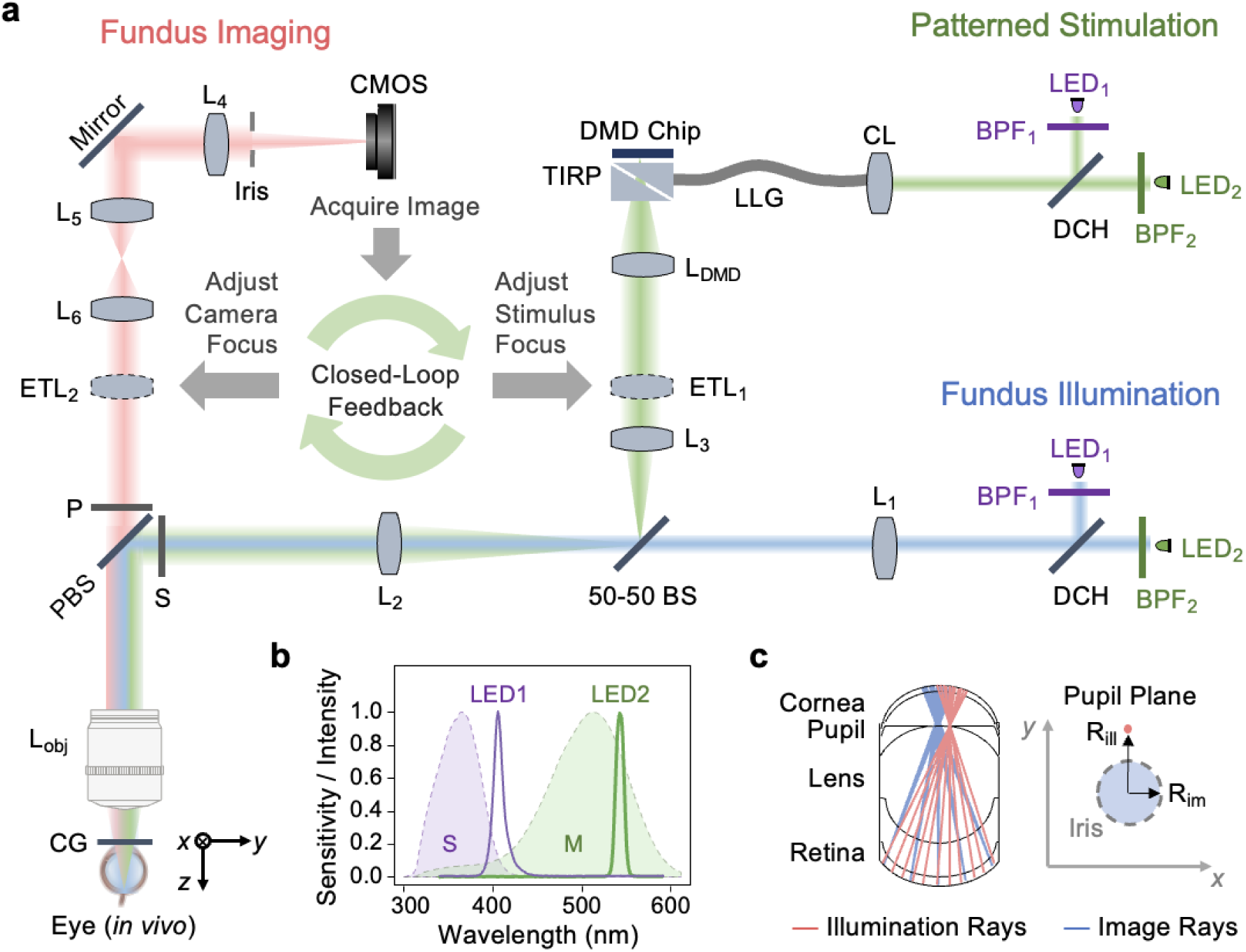
Experimental system. (a) Optical schematics of the experimental system, divided into three optical paths: fundus illumination (blue), patterned stimulation (green), and fundus imaging (red). CG: cover glass, PBS: polarizing beamsplitter, ETL: electronically tunable lens, CMOS: complementary metal-oxide semiconductor camera, BS: beamsplitter, TIRP: total internal reflection prism, DMD: digital micromirror device, LLG: liquid light guide, DCH: dichroic mirror, LED: light-emitting diode, BPF: bandpass filter. (b) Spectral profiles of mouse S- an M-cone sensitivities overlaid with the emission spectra of the LEDs. (c) Angular separation between the illumination and imaging rays at the pupil plane. R_ill_: lateral offset of the illumination source, R_im_: radius of the image beam.

The fundus illumination path (colored in blue) provides the light required for retinal imaging. We optically conjugated two light-emitting diodes (LEDs) to the pupil plane, following the principles of Köhler illumination to minimize specular reflections while achieving uniform retinal irradiance [24]. Light from each LED passes through lenses L_1_, L_2_, and L_obj_ before entering the eye. To mitigate chromatic aberration, the same LEDs used for patterned stimulation are used for fundus illumination. Bandpass filters (±5 nm, BPF_1_ and BPF_2_) were placed in front of LED_1_ and LED_2_, respectively, to ensure spectral specificity, and the two channels were combined using a dichroic mirror (DCH).

The patterned stimulation path (colored in green) originates from two LEDs (LED_1_ and LED_2_), whose wavelengths were chosen to activate distinct cone subtypes in the mouse retina as shown in **Fig. 1(b)**. LED_1_ (centered at 415 nm after BPF_1_) was chosen as the primary alignment source because it provides balanced activation of S- and M-cones in the mouse retina [25]. LED_2_ (centered at 550 nm after BPF_2_) preferentially activates M-cones. A dedicated S-cone-activating source was not included because dorsal M-cones co-express opsins that remain sensitive to shorter wavelengths [26]. Light from both sources is directed to the DMD to generate the desired patterns, which are then relayed to the retina through a pair of telescopic systems composed of lenses L_obj_, L_2_, L_3_, and L_DMD_. We placed an ETL (ETL_1_) at a pupil conjugate plane to adjust the stimulus focus.

Patterned stimuli were referenced to a retinal irradiance of 1 µW/mm^2^ and scaled by the photoreceptor sensitivity function *S(Ż)* to account for wavelength-dependent activation [25]. The standardized radiant power, *P_c_* (in µW), to be delivered through the pupil is given by

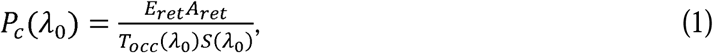

where *E*_ret_ is the retinal irradiance (in µW/mm^2^), *A* is the retinal area being stimulated (in mm^2^), *T*_Occ_ *(λ*_0_) is the ocular transmission, and *S(λ*_0_) is the sensitivity at the specified wavelength. Using literature values for adult BALB/c mice [25, 27], the objective-output power was calibrated to 56.8 µW (LED_1_) and 11.4 µW (LED_2_) under full-field conditions, in which all micromirrors are engaged as shown in **Supplementary Methods S1**.

The fundus imaging path (colored in red) provides real-time visualization of projected patterns on the retina, enabling verification of stimulus location and focus optimization. A monochrome complementary metal-oxide semiconductor (CMOS) camera was conjugated to the retina via a telescopic relay comprising lenses L_4_, L_5_, L_6_, and L_obj_. A second ETL (ETL_2_) was placed at the pupil-conjugate plane closest to the camera to address chromatic defocus when switching to a different stimulus wavelength without having to reposition the animal. Because specular reflections from the fundus illumination and patterned stimulation paths can oversaturate retinal features, a polarizing beam splitter (PBS) and a set of cross-polarizers (S and P) were added at the point of separation [28]. S-polarized light from the LEDs is directed towards the eye by the PBS, and the backscattered light becomes partially depolarized upon interacting with the retina. On the return path, the analyzing polarizer (P) allows only the orthogonally polarized component of the reflected light to reach the CMOS. The extinction ratios were 673:1 and 702:1 for LED_1_ and LED_2_, respectively.

Additional suppression of back-reflected light was achieved through angular separation between the illumination and image beams at the pupil plane, as shown in **Fig. 1(c)** [29]. The fundus illumination LED (depicted as a pink point source) was laterally displaced from the optical axis, with R_ill_ denoting its offset in the pupil plane. The fundus imaging path is limited by a circular aperture (shown as a blue disc) of radius R_im_, centered on the optical axis and defined by an adjustable iris placed at the nearest pupil-conjugate plane to the CMOS. Empirically, we found that a ratio R_ill_/R_im_ ≈ 1.5 provided optimal performance [**Fig. S2**]. Lower ratios increase overlap between the illumination and imaging paths, leading to stronger corneal reflections, whereas higher ratios reduce the amount of light entering the pupil, thereby decreasing the retinal signal.

### 2.2 Projection performance in an ideal conjugate plane

To evaluate projection quality under ideal optical conditions, we generated a 10° slanted edge using the DMD and imaged it at the retinal conjugate plane using a secondary CMOS camera. Direct imaging of the DMD allows us to isolate the intrinsic optical performance of the pattern-stimulation path from any influence of the mouse eye. For quantitative assessment, we divided the slanted-edge image into nine horizontal blocks and computed an edge-spread function (ESF) for each block using a modified version of ISO 12233, as described in Section 4.2 [30]. **Fig. 2(a)** shows the averaged ESF (red) overlaid on the individual ESF profiles (gray), with the inset showing the imaged slanted edge.

**Figure 2.**
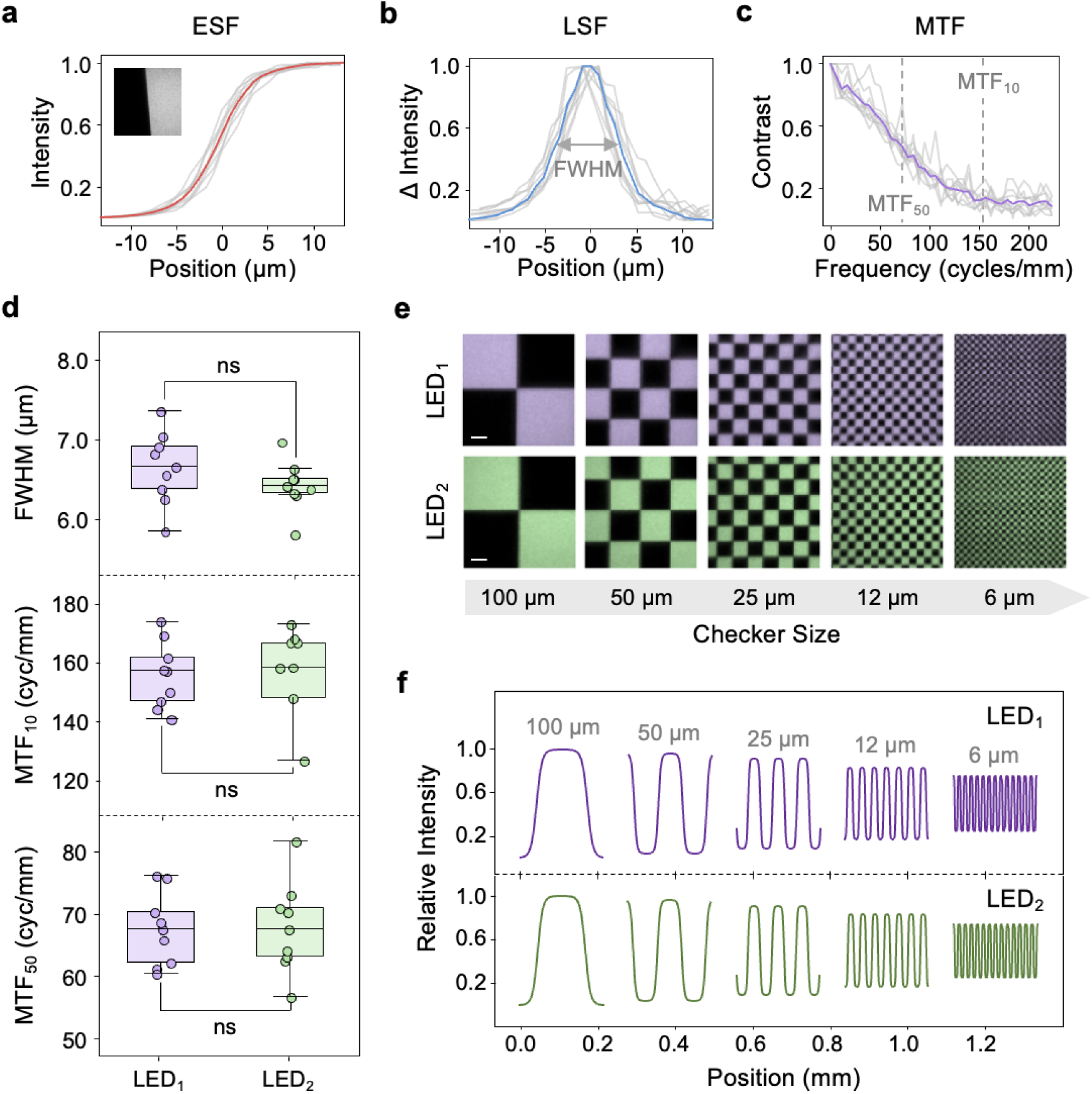
Quantification of stimulus quality. (a) ESFs obtained by imaging a 10°-tilted edge projected from the DMD onto a camera positioned at the retinal conjugate plane using LED_1_. The mean ESF is colored in red. (b) Corresponding LSFs by taking the first derivatives of the ESFs, with the mean LSF colored in blue. The FWHM of the LSF was used as a direct measurement of the stimulus lateral resolution. (c) MTFs obtained from the Fourier transform of the LSFs, with the mean MTF colored in purple. MTF_50_ and MTF_10_ denote the spatial frequencies at which contrast decreases to 50% and 10%, respectively. (d) Summary of resolution and contrast metrics for LED_1_ and LED_2_ following tunable lens correction. No significant differences were observed. (e) Projected checkerboard patterns with decreasing feature sizes. Scale bars: 25 µm. (f) Corresponding intensity profiles demonstratin preserved periodicity and contrast across spatial frequencies. Modulation remains detectable at a 6 µm checker size, consistent with the measured MTF limits.

Lateral resolution was quantified by differentiating the ESFs to obtain the corresponding line spread functions (LSFs) and measuring their full widths at half maximum (FWHMs) [**Fig. 2(b)**]. Contrast transfer was evaluated using the modulation transfer function (MTF) [**Fig. 2(c)**], computed as the Fourier transform of the LSFs [31]. Each MTF was normalized to unity at zero frequency, and the analysis was restricted to positive spatial frequencies. The frequencies at which the modulation dropped to 50% (MTF_50_) and 10% (MTF_10_) of the zero-frequency value were extracted as summary metrics. The same process was repeated with LED_2_ after ETL correction [**Fig. S3**]. No significant differences were observed between the two LEDs in FWHM, MTF_10_, or MTF_50_ values. In all cases, the average lateral resolution remained below 6.7 µm, with MTF_10_ exceeding 150 cycles/mm, and MTF_50_ exceeding 65 cycles/mm [**Fig. 2(d)**]. The complete results are provided in **Table S1**.

To validate these findings, we assessed lateral resolution and contrast in the spatial domain by projecting DMD-generated checkerboard patterns with feature sizes ranging from 100 µm down to 6 µm, the latter corresponding to approximately twice the linear dimension of a single CMOS pixel. Representative images of the projected checkerboards for LED_1_ and LED_2_ are shown in **Fig. 2(e)**. Furthermore, one-dimensional intensity modulations were extracted from these patterns by taking each row, aligning the checker peaks to a common reference, and averaging across rows. The averaged profiles were then fit to a flattened sine function as

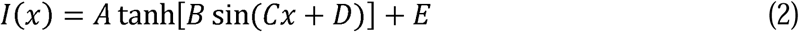

using nonlinear least squares. The resulting modulations [**Fig. 2(f)**] represent the spatial periodicity of the respective checkerboards, with peak-to-trough amplitudes serving as direct measures of contrast [**Table S2**]. At the smallest checker size, corresponding to a spatial frequency of 83 cycles/mm, the contrast is approximately equal to the MTF_50_, placing it in a regime where high-frequency components begin to be attenuated, though core structures remain well resolved.

### 2.3 Characterization of in vivo stimulus quality

*In vivo* stimulus quality is influenced by additional factors such as ocular aberrations and anatomy [8]. To understand these effects, we analyzed the patterned stimulation path using ray-tracing (Ansys Zemax OpticStudio) and an optical mouse-eye model [**Fig. S4**] [11]. First, crystalline lens-induced chromatic aberration was evaluated by quantifying the shift in the stimulus focal plane between the two LEDs. The resulting focal shift was 230 µm [**Fig. 3(a)**], comparable to the full axial span of the BALB/c mouse retina [32]. This prediction was experimentally validated by first optimizing focus with LED_1_ using the Tenengrad metric (see Section 4.4), then switching to LED_2_ and axially translating the mice (*n* = 5) until focus was re-established. The measured displacement was 213 ± 43 µm (mean ± SD) [**Fig. 3(b)**], consistent with the ray-tracing analysis.

**Figure 3.**
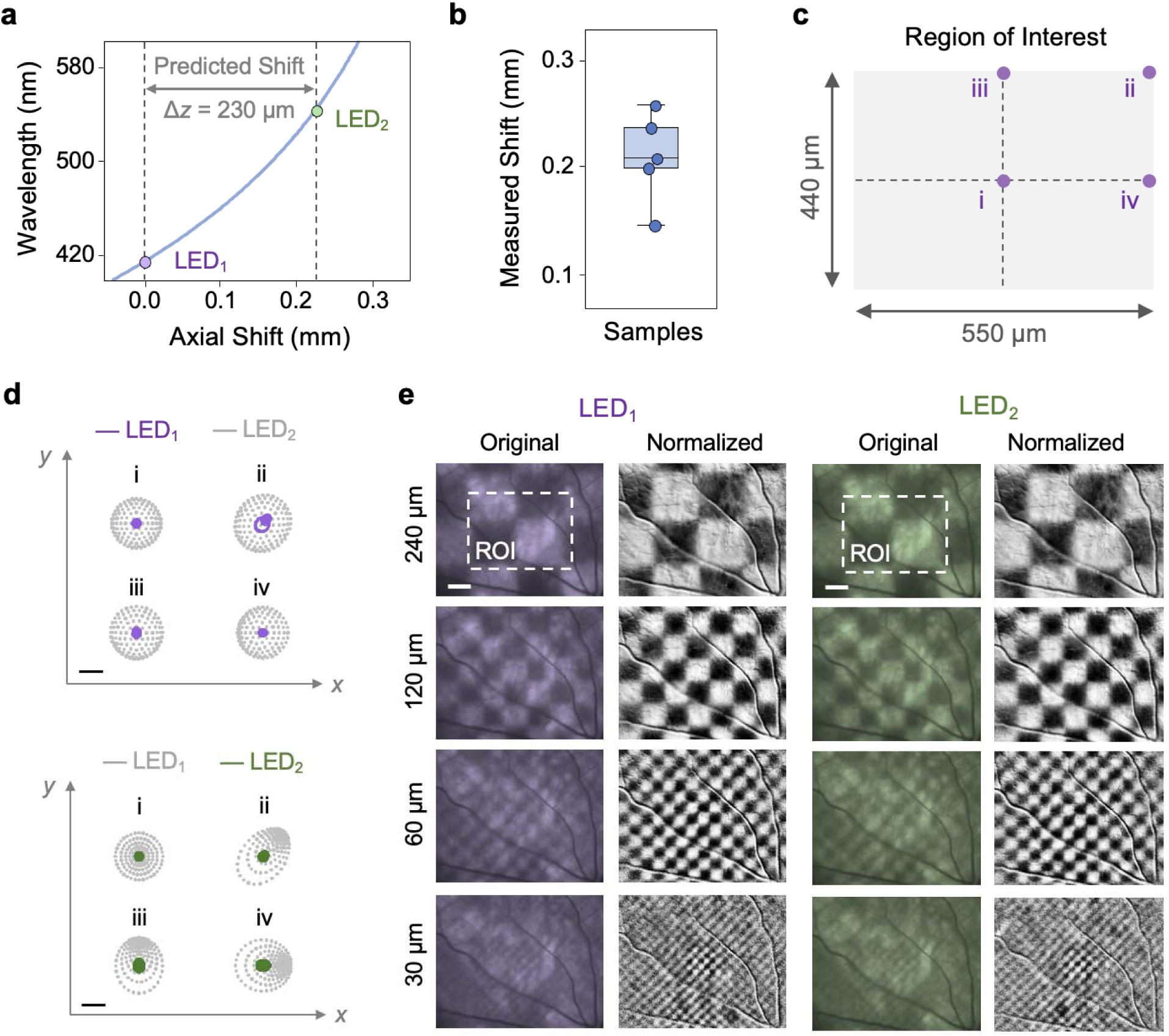
Characterization of the optical properties of the mouse eye. (a) Ray-tracing simulations showing the axial focal shift as a function of wavelength. The focal positions of LED_1_ and LED_2_ are highlighted and separated by Δ*z* ≈ 230 µm, which is comparable to the retinal thickness. (b) Experimentally measured axial shift (213 ± 43 µm) obtained using Tenengrad-based focus optimization across samples. (c) A 550 × 400 µm^2^ ROI used for analysis, with evaluation points at the (i) center, (ii) corner, (iii) midpoint of the long-axis edge, and (iv) midpoint of the short-axis edge. (d) Ray-tracing spot diagrams at the four locations (i-iv) within the ROI, showing reduced spot size with cover glass application and chromatic defocus correction. Colored spots denote the active wavelength (purple: LED_1_; green: LED_2_), while gray spots indicate the opposite wavelength under identical conditions. Residual deviations at the periphery arise from higher-order aberrations, including coma and field curvature. Scale bars: 20 µm. (e) *In vivo* fundus images of projected stimuli for LED_1_ and LED_2_ with decreasing feature sizes. The ROI is highlighted by the dashed boxes. Corresponding normalized images, obtained by division with a uniform illumination reference, enhance contrast for visualization. ROI: region of interest. Scale bars: 120 µm.

Although chromatic defocus can be corrected by the ETL, the high curvature of the mouse eye (*x* ≈ 0.7 mm^-1^) introduces additional field-dependent aberrations that can degrade stimulus quality [10]. A simple approach to limit this type of aberration is to apply a cover glass to the cornea [33, 34], which reduces the focal power by creating a flattened geometry, as shown by the cross-sectional optical coherence tomography image in **Fig. S5**. Another strategy is to reduce the stimulation FOV or restrict analysis to a region of interest (ROI) centered on the optical axis. We quantified these effects using ray-tracing analyses to characterize four points within a 550 × 400 µm^2^ ROI: the (i) center, (ii) corner, (iii) midpoint of the long edge, and (iv) midpoint of the short edge [**Fig. 3(c)**]. Spot diagrams and point spread functions (PSFs) were obtained at each location before [**Figs. S6** and **S7**, **Table S3**] and after [**Figs. 3(d)** and **S8**] cover glass application, where all root-mean-square (RMS) spot radii remained under 5.0 µm within the ROI [**Table S4**].

To experimentally validate the ray-tracing analyses, we projected checkerboard patterns onto mouse retinas and imaged the pattern-illuminated retinas *in vivo*. We optimized the fundus imaging and stimulation patterns using the Tenengrad metric, starting from a checker size of 240 µm [**Fig. 3(e)**]. The pattern-illuminated fundus images were normalized by a reference fundus image acquired under uniform illumination to enhance the visualization of the patterns. We also performed a frequency sweep at 0.7 octaves per second, starting with a checker size of 500 µm and continuing until the checkerboard patterns became unresolved [**Supplementary Movie S1**]. Patterns were clearly resolved until checkerboard sizes approached the size of the primary blood vessels of the superficial vascular plexus (around 30 µm in diameter) [35], at which point hemoglobin absorption introduced localized attenuation. Additional projections of symbols and geometric patterns further verified the accuracy of pattern formation [**Fig. S9** and **Supplementary Movie S2**].

## 3 Discussion and Conclusion

In this study, we developed and characterized a closed-loop optical platform that enables patterned stimulation of the mouse retina *in vivo* while directly verifying both stimulus location and focus. Rather than relying on intact optics, as in conventional external displays or infinity-focused projection systems [13–16], our approach continuously visualizes the projected stimulus on the retina and optimizes its focus using a gradient-based feedback metric, establishing a reproducible framework for delivering patterned stimuli across subjects and experimental sessions. This closed-loop strategy is further supported by ETLs that permit rapid switching between stimulation wavelengths while maintaining focus. Optical characterization demonstrated lateral resolutions below 6.7 µm in the ideal conjugate plane, while ray-tracing simulations predicted RMS spot radii below 5.0 µm within the central ROI. Together, these results establish the optical performance necessary for cellular-scale retinal stimulation. Because this resolution is comparable to the average spacing between cones in the mouse retina, approximated to be 9 µm [**Supplementary Methods S2**] [36], patterned optical stimuli can be confined to small groups of cells, rather than broad retinal regions. This capability allows retinal circuitry to be interrogated at cellular scales, a level of precision previously unattainable with conventional *in vivo* patterned stimulation methods.

Such control over stimulus pattern formation *in vivo* has important implications for vision research. Precise retinal patterning enhances the interpretability of assays, such as receptive-field mapping [37, 38] and pattern electroretinography [39], which depend on spatial contrast. Although we previously achieved single-cell-resolution Ca^2+^ imaging of RGCs in live animals [7], the lack of spatially distributed and feature-selective visual stimuli, including direction, orientation, temporal, contrast, and chromatic modulation, limits its application. Our system addresses this gap by enabling closed-loop image-guided delivery of diverse visual patterns to the mouse retina. These include size-adjustable spots and white noise for receptive-field mapping [4], moving bars for direction and orientation selectivity [40], and wavelength-specific patterns for chromatic responses [41]. This flexibility supports functional interrogation across multiple stages of the visual pathway, including the retina, the lateral geniculate nucleus, the superior colliculus, and the visual cortex [42]. It also normalizes longitudinal studies, in which age, disease, or genetic factors may alter the focusing power of the intact mouse eye over time [7, 43]. By verifying retinal stimulus placement while minimizing variability in optical quality, this approach allows observed differences to be more reliably attributed to retinal physiology rather than differences in optical focus or retinal alignment.

Although the ideal plane characterization demonstrated near-diffraction-limited performance, image quality was reduced under *in vivo* conditions. This degradation can be attributed to several factors, including the round-trip nature of the measurement, field-dependent defocus introduced by the retinal curvature, absorption and scattering from the retinal vasculature, and higher-order aberrations from the intact mouse eye [8]. Importantly, the projected stimulus is delivered to the retina during the forward optical path, whereas the fundus camera measures only the attenuated light returning from the retina. The deeper layers of the retina, such as the pigment epithelium and choroid [30], are also specialized to absorb rather than reflect light, further reducing the returning signal. Consequently, the measured contrast and resolution represent conservative estimates of the stimulus quality experienced by the photoreceptors. Despite these limitations, retinal features as small as 30 µm remained discernible in the central region of interest, supporting reliable closed-loop alignment. Because these images are formed from the attenuated return signal rather than the forward-propagating stimulus, the optical quality delivered to the photoreceptors is expected to exceed that observed by the fundus camera.

The limitations of the present study suggest several directions for future improvement, particularly in addressing higher-order, field-dependent aberrations that are difficult to correct solely by ETLs. Although we have shown that restricting the stimulus FOV is effective in minimizing these effects, integrating adaptive optics (AO) can extend this performance across the full FOV [44]. By dynamically correcting spatially varying wavefront distortions, AO enables more uniform resolution across the retinal field, including peripheral regions. Both wavefront-sensing AO, which uses dedicated sensors to measure aberrations [45], and sensorless AO, which infers corrections by optimizing image-based metrics [46], may be compatible with the current architecture. Another direction is to further integrate stimulus delivery with functional readouts, such as calcium signals [47] or electrophysiological recordings [48], to enable dynamic adaptation of stimulus parameters. This would shift the system from fixed-pattern delivery to one that is guided by ongoing neural activity. Additionally, vascular landmarks could be leveraged for real-time motion correction, enabling more precise registration of the stimulus to the retina.

In summary, our work establishes a methodological framework for patterned retinal stimulation in which stimulus formation at the retinal plane is quantitatively verified and corrected in real-time. By coupling high-resolution DMD patterning with closed-loop focus optimization, the system addresses a longstanding limitation of *in vivo* retinal stimulation by directly verifying where patterned stimuli are delivered on the retina while correcting focus in real time. Our approach enables reproducible, cellular-scale delivery of structured visual stimuli while supporting precise multi-wavelength retinal stimulation through dynamic correction of chromatic aberration. Together, these capabilities provide a foundation for future studies of disease progression, retinal circuitry, color processing, and optical manipulation of the visual pathway.

## 4 Materials and Methods

### 4.1 Optical components

Fundus illumination was provided by LED_1_ (Thorlabs M415L4, *Ż*_0_ = 415 nm) and LED_2_ (Thorlabs M565L3, *Ż*_0_ = 565 nm). Each source was spectrally filtered using 10 nm bandpass filters (FBH410-10 for LED_1_; FBH550-10 for LED_2_) before being combined onto a common optical axis with a long-pass dichroic mirror (Thorlabs DMLP505R). The combined beam was relayed to the pupil plane through a telescopic system consisting of L_1_ (*f* = 42.5 mm, Edmund Optics #47-349), L_2_ (*f* = 250 mm, Edmund Optics #88-736), and L_obj_ (Thorlabs TL10X-2P).

The patterned stimulation path uses an identical set of LEDs, bandpass filters, and a dichroic mirror. Spatial patterns were generated using a DMD-based light engine (EKB Technologies DPM-E4500MKIIFC-OX) incorporating a TIR prism and condensing lens (L_DMD_), mounted on *xy*- (Thorlabs XYT1) and *z*-translation stages (Thorlabs MVS05). The LEDs were coupled to the light engine via a liquid light guide (Thorlabs LLG05-4H), enabling flexible positioning of the source. The DMD chip was relayed through L_DMD_, L_3_ (*f* = 37.5 mm, Edmund Optics #88-726), L_2_, and L_obj_, resulting in a 4× demagnification. ETL_1_ (Optotune EL-16-40-TC-VIS-20D) was placed between L_3_ and L_obj_ for focus adjustment, and a 50:50 beamsplitter (Thorlabs BSW26R) was used to switch between the patterned stimulation and illumination pathways.

Fundus imaging was achieved using a monochrome CMOS camera (Teledyne FLIR BFS-U3-16S2M-BD2), which was conjugated to the retina using telescopic elements L_4_ (*f* = 37.5 mm, Edmund Optics #88-726), L_5_ (*f* = 62.5 mm, Edmund Optics #88-729), L_6_ (*f* = 75 mm, Edmund Optics #88-730), and L_obj_, providing a 4.5× magnification. ETL_2_ (Optotune EL-16-40-TC-VIS-20D) was placed between L_6_ and L_obj_ for stimulus focus correction. Polarization-based reflection suppression was implemented using a wire grid broadband PBS (Edmund Optics #48-545) and linear cross-polarizers (Thorlabs WP25M-UB1). An iris diaphragm (Thorlabs SM1D12) was placed at a pupil-conjugate plane in front of the camera to define the imaging beam radius.

The system was implemented in a 30 mm cage configuration consisting of cage rods (Thorlabs ER1-ER8), cage plates (Thorlabs CP33), lens tubes (Thorlabs SM1L10), and beamsplitter cubes (Thorlabs CM1-DCH). Optical posts (Thorlabs TR3-TR4) were mounted in post holders (Thorlabs PH3-PH4) and secured to base plates (Thorlabs BE1). The objective lens was integrated using an SM1-thread adapter (Thorlabs M1M32S). The numbered lenses (L_1_-L_6_) were arranged in paired plano-convex relay configurations, with their planar surfaces facing outward and curved surfaces facing inward. This symmetric pairing yields an effective focal length that is half that of a single element and reduces spherical aberration relative to a single-lens relay.

### 4.2 Modified slanted-edge method

A modified slanted-edge method was used to acquire the ESF, overcoming the limitations of pixel-limited sampling. First, a 10° binary edge was projected onto the target of interest, and the resulting image */(x, y)* was divided horizontally into 9 equal-sized blocks to assess field-dependent variations.

Edge localization was then carried out within each block by computing the *x*-direction derivative *p*_r_*(x)* along each row *y*_r_ using a first-order finite difference. The edge position 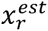 was initially estimated as the location of the maximum gradient magnitude *arg max*_x_ *|p*_r_*(x)|*. To achieve subpixel precision, this estimate was refined by fitting a quadratic to the neighboring gradient samples 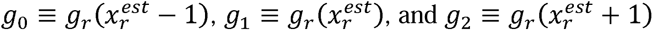, yielding

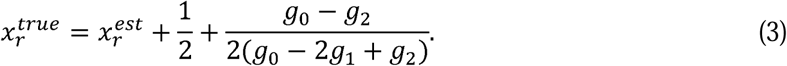

This procedure produces a set of edge points 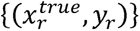 describing the edge within the block. A straight line was then fit to these edge points using least squares and expressed in the implicit form *ax + by + c — 0*. The perpendicular distance of each pixel *(x_i_, y_i_(* to the fitted edge was computed as

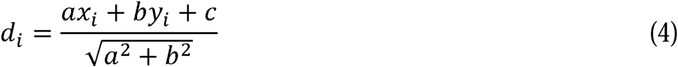

and converted to physical units via 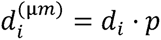, where *p* is the effective pixel size in micrometers.

Finally, all pixels within the block were flattened into a set of samples *(d*_i_*, /*_i_), where *I_i_ — /(x_i_, y_i_(*, and the ESF was constructed by binning along the distance axis. The distance values were discretized with a binning factor of 4, and the ESF at each bin center *d_j_* was computed as the average intensity of all pixels whose distances fell within that bin:

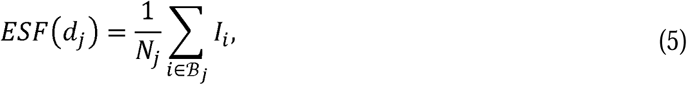

where *B*_j_ denotes the set of pixels in bin *J*, and *N*_j_ is the number of samples in that bin. Differentiating this equation yields the LSF, and applying a Fourier transform to the LSF yields the MTF. These functions were computed independently across the 9 blocks, and their resulting metrics (e.g., FWHM of the LSF, MTF_10_, and MTF_50_) were summarized by their means and standard deviations to assess average performance and field-dependent variation.

### 4.3 Mouse preparation and handling

Before experiments, adult BALB/c mice (The Jackson Laboratory #000651) were anesthetized by intraperitoneal injection of Ketamine (11.45 mg/mL) and Xylazine (1.70 mg/mL) at 10 mL/kg. Pupil dilation was achieved with 1-2 drops of 1% topical tropicamide, and artificial tears were applied to maintain corneal hydration. Animals were secured in a custom holder, equipped with a bite bar for stabilization. Before imaging, a cover glass (Fisher Scientific #12-541-039) was gently placed in contact with the cornea to reduce optical power and mitigate higher-order aberrations that could otherwise degrade stimulus quality [33, 34]. Body temperature was maintained between 35.5 and 38.0 °C using a heat lamp, and physiological status was continuously monitored to ensure stable anesthetic depth. All procedures in this study were performed in accordance with a protocol approved by the Northwestern University Animal Care and Use Committee (#IS00023786).

### 4.4 Closed-loop retinal stimulus localization and focus adjustment

Accurate retinal conjugation is essential for proper formation of patterned stimuli. Because the superficial vascular plexus provides excellent contrast at the lower end of the visible spectrum, the blood vessels in this region were used as a reference for focus. To ensure this corresponds to the photoreceptor layer, where phototransduction begins, the depth of focus (DOF) of the stimulus pathway was set to span from the vessels to the photoreceptor outer segments, which is around 180 µm for mice [**Fig. 4(a)** and **Supplementary Methods S3**] [49]. This configuration also maintains a diffraction-limited lateral resolution below 7 µm. Retinal alignment was performed by placing the mouse on a translation stage with one eye facing the objective lens and adjusting the degrees of freedom (3 axial, 3 angular) until the optic nerve head (ONH) became visible. The ONH was positioned at the corner of the FOV to preserve regions containing higher densities of photoreceptors.

**Figure 4.**
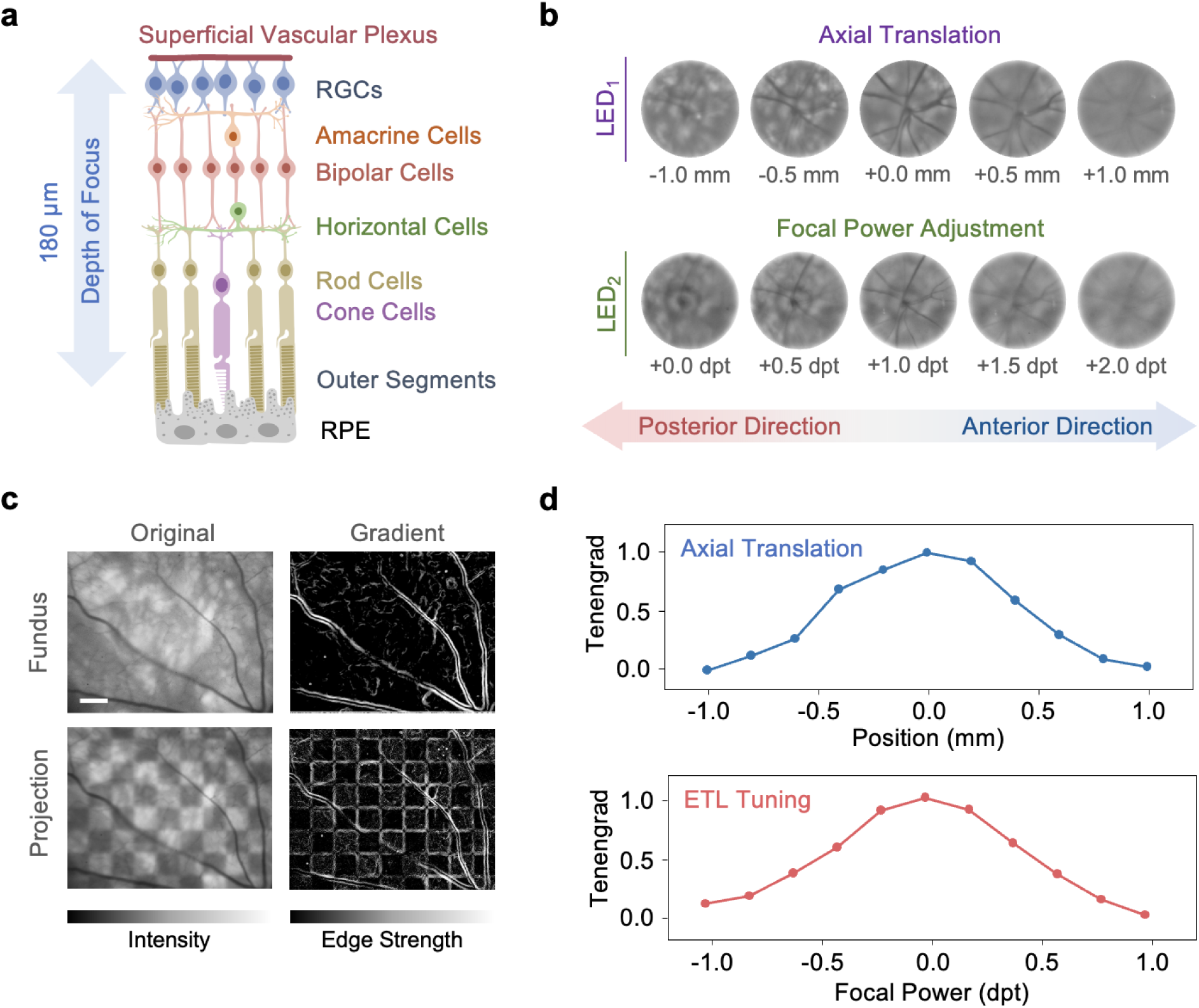
*In vivo* alignment and focus optimization. (a) Schematic of retinal layers illustrating the required depth of focus, which spans from the superficial vascular plexus to the photoreceptor outer segments (∼180 µm). This ensures that vascular contrast can guide alignment while maintaining focus at the stimulation plane. (b) Focus optimization for imaging and stimulation. For LED_1_, the optimal focus was identified by axially translating the mouse to maximize vessel sharpness. For LED_2_, chromatic defocus was corrected by adjusting the ETL. Dpt: diopters. (c) Representative fundus images under uniform illumination (top) and patterned projection (bottom), shown alongside their corresponding gradient magnitude images used for focus evaluation. Scale bar: 120 µm. (d) Tenengrad focus metric as a function of axial position (top) and ETL focal power (bottom). The metric, defined as the normalized mean squared gradient magnitude, is maximized at optimal focus.

Optimal focus of the fundus imaging was first identified by performing an axial sweep of the translation stage using the primary illumination source (LED_1_) [**Figs. 4(b)** and **S10**]. Switching to a different wavelength introduces chromatic defocus, which was compensated for by adjusting ETL_2_. Once the camera focus was established, a checkerboard pattern of the same wavelength was projected, and ETL_1_ was swept to optimize the stimulation patterns.

For both processes, image sharpness was quantified in real time using the Tenengrad metric [50]. This was computed by first evaluating the image gradients *G*_x_ and *G*_y_ with a Sobel operator (kernel size = 9) in the horizontal and vertical directions, respectively [51]. This allowed us to reconstruct a gradient map,

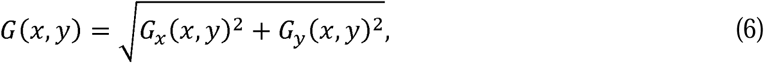

as illustrated in **Fig. 4(c)** for uniformly illuminated and patterned-illuminated fundus images. The Tenengrad metric *T* was then calculated as the mean squared gradient over the image:

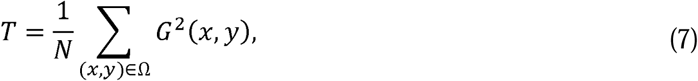

where *Ω* denotes the set of valid pixels and *N* is the total number of pixels. Maximizing *T* over the range of axial translation or ETL tuning signifies that the image is in focus [**Fig. 4(d)**]. A flowchart summarizing the retinal alignment process is shown in **Fig. S11**.

Lastly, patterned stimulation operates in the photopic regime and is therefore primarily cone-driven. During alignment, illumination and stimulus powers were kept below the limits specified in Section 2.1 to avoid overstimulation. Although cones can recover within a few hundred milliseconds after stimulation, a 1-minute dark interval was introduced to ensure a stable retinal baseline prior to any patterned stimulation experiments [52].

### 4.5 Optical Power and Spectral Characterization

Optical power exiting the objective lens was measured using a photodiode power meter (Thorlabs PM400K1). LED drive currents were adjusted to achieve the target retinal irradiance described in Section 2.1. The spectral output of each LED, after passing through its respective bandpass filter, was measured using a fiber-coupled spectrometer (Ocean Optics Maya LSL) to verify the center wavelength and bandwidth.

### 4.6 Statistical analysis

Statistical analysis was conducted to assess whether the two LEDs differed significantly in lateral resolution and contrast, using metrics such as FWHM, MTF_10_, and MTF_50_. Values lying more than three standard deviations from the group mean were excluded as outliers. Equality of variances between groups was first assessed using Levene’s test, which determined the appropriate parametric comparison. If variances were not significantly different (*p* ≥ *ce*), a two-tailed Student’s *t*-test was applied. Conversely, if variances were unequal (*p* < *ce*), Welch’s *t*-test was used instead. In all cases, a significance threshold of *ce* = 0.05 was used. All statistical analyses were conducted in Python using the SciPy library.

## Acknowledgements

We thank Prof. Sava Sakadzic and Prof. Wei Wei for the insightful discussions regarding the optical design and mouse setup.

## Data availability

Data underlying the results presented in this paper are not publicly available at this time but may be obtained from the authors upon reasonable request.

## References

1. J. Chen, R. Fang, X. Liu et al., “Optical strategies for in vivo retinal ganglion cell imaging,” Med X 3, 21 (2025).

2. E. S. Ruthazer, and C. D. Aizenman, “Learning to see: patterned visual activity and the development of visual function,” Trends Neurosci 33, 183–192 (2010).

3. D. Gupta, W. Mlynarski, A. Sumser et al., “Panoramic visual statistics shape retina-wide organization of receptive fields,” Nat Neurosci 26, 606–614 (2023).

4. T. Baden, P. Berens, K. Franke et al., “The functional diversity of retinal ganglion cells in the mouse,” Nature 529, 345–350 (2016).

5. G. T. Prusky, N. M. Alam, S. Beekman et al., “Rapid quantification of adult and developing mouse spatial vision using a virtual optomotor system,” Invest Ophthalmol Vis Sci 45, 4611–4616 (2004).

6. T. H. Chou, J. Toft-Nielsen, and V. Porciatti, “Adaptation of retinal ganglion cell function during flickering light in the mouse,” Sci Rep 9, 18396 (2019).

7. L. Li, X. Feng, F. Fang, et al., “Longitudinal in vivo Ca(2+) imaging reveals dynamic activity changes of diseased retinal ganglion cells at the single-cell level,” Proc Natl Acad Sci U S A 119, e2206829119 (2022).

8. Y. Geng, A. Dubra, L. Yin et al., “Adaptive optics retinal imaging in the living mouse eye,” Biomed Opt Express 3, 715–734 (2012).

9. Y. Geng, L. A. Schery, R. Sharma et al., “Optical properties of the mouse eye,” Biomed Opt Express 2, 717–738 (2011).

10. C. Schmucker, and F. Schaeffel, “A paraxial schematic eye model for the growing C57BL/6 mouse,” Vision Res 44, 1857–1867 (2004).

11. S. Remtulla, and P. E. Hallett, “A schematic eye for the mouse, and comparisons with the rat,” Vision Res 25, 21–31 (1985).

12. M. T. Pardue, R. A. Stone, and P. M. Iuvone, “Investigating mechanisms of myopia in mice,” Exp Eye Res 114, 96–105 (2013).

13. X. J. Chen, M. J. Rasch, G. Chen et al., “Binocular input coincidence mediates critical period plasticity in the mouse primary visual cortex,” J Neurosci 34, 2940–2955 (2014).

14. D. R. Muir, M. M. Roth, F. Helmchen et al., “Model-based analysis of pattern motion processing in mouse primary visual cortex,” Front Neural Circuits 9, 38 (2015).

15. H. Tabata, N. Shimizu, Y. Wada et al., “Initiation of the optokinetic response (OKR) in mice,” J Vis 10, 13 11–17 (2010).

16. T. Arens-Arad, N. Farah, S. Ben-Yaish et al., “Head mounted DMD based projection system for natural and prosthetic visual stimulation in freely moving rats,” Sci Rep 6, 34873 (2016).

17. J. B. Demb, L. Haarsma, M. A. Freed et al., “Functional circuitry of the retinal ganglion cell’s nonlinear receptive field,” J Neurosci 19, 9756–9767 (1999).

18. L. Y. Yu, and S. You, “High-fidelity and high-speed wavefront shaping by leveraging complex media,” Sci Adv 10, eadn2846 (2024).

19. M. J. Deng, Y. Y. Zhao, Z. X. Liang et al., “Maximizing energy utilization in DMD-based projection lithography,” Opt Express 30, 4692–4705 (2022).

20. Y. V. Wang, M. Weick, and J. B. Demb, “Spectral and temporal sensitivity of cone-mediated responses in mouse retinal ganglion cells,” J Neurosci 31, 7670–7681 (2011).

21. S. Yang, X. Luo, G. Xiong et al., “The electroretinogram of Mongolian gerbil (Meriones unguiculatus): comparison to mouse,” Neurosci Lett 589, 7–12 (2015).

22. K. Farrow, and R. H. Masland, “Physiological clustering of visual channels in the mouse retina,” J Neurophysiol 105, 1516–1530 (2011).

23. L. Chen, M. Ghilardi, J. J. C. Busfield et al., “Electrically Tunable Lenses: A Review,” Front Robot AI 8, 678046 (2021).

24. M. Saxena, G. Eluru, and S. S. Gorthi, “Structured illumination microscopy,” Advances in Optics and Photonics 7 (2015).

25. L. Abballe, and H. Asari, “Natural image statistics for mouse vision,” PLoS One 17, e0262763 (2022).

26. T. Baden, T. Euler, and P. Berens, “Understanding the retinal basis of vision across species,” Nat Rev Neurosci 21, 5–20 (2020).

27. J. T. Henriksson, J. P. Bergmanson, and J. E. Walsh, “Ultraviolet radiation transmittance of the mouse eye and its individual media components,” Exp Eye Res 90, 382–387 (2010).

28. D. W. Palmer, T. Coppin, K. Rana et al., “Glare-free retinal imaging using a portable light field fundus camera,” Biomed Opt Express 9, 3178–3192 (2018).

29. E. DeHoog, and J. Schwiegerling, “Fundus camera systems: a comparative analysis,” Appl Opt 48, 221–228 (2009).

30. X. Xie, H. Fan, H. Wang, et al., “Error of the slanted edge method for measuring the modulation transfer function of imaging systems,” Appl Opt 57, B83–B91 (2018).

31. A. S. Chawla, H. Roehrig, J. J. Rodriguez et al., “Determining the MTF of medical imaging displays using edge techniques,” J Digit Imaging 18, 296–310 (2005).

32. C. Dysli, V. Enzmann, R. Sznitman et al., “Quantitative Analysis of Mouse Retinal Layers Using Automated Segmentation of Spectral Domain Optical Coherence Tomography Images,” Transl Vis Sci Technol 4, 9 (2015).

33. B. E. Cohan, A. C. Pearch, P. T. Jokelainen et al., “Optic disc imaging in conscious rats and mice,” Invest Ophthalmol Vis Sci 44, 160–163 (2003).

34. X. Liu, C. H. Wang, C. Dai et al., “Effect of contact lens on optical coherence tomography imaging of rodent retina,” Curr Eye Res 38, 1235–1240 (2013).

35. F. Shang, and J. Schallek, “Characterization of the Retinal Circulation of the Mouse,” Invest Ophthalmol Vis Sci 65, 3 (2024).

36. C. J. Jeon, E. Strettoi, and R. H. Masland, “The major cell populations of the mouse retina,” J Neurosci 18, 8936–8946 (1998).

37. K. Franke, A. Maia Chagas, Z. Zhao et al., “An arbitrary-spectrum spatial visual stimulator for vision research,” Elife 8 (2019).

38. J. Freeman, G. D. Field, P. H. Li et al., “Mapping nonlinear receptive field structure in primate retina at single cone resolution,” Elife 4 (2015).

39. G. E. Holder, “Pattern electroretinography (PERG) and an integrated approach to visual pathway diagnosis,” Prog Retin Eye Res 20, 531–561 (2001).

40. D. I. Vaney, B. Sivyer, and W. R. Taylor, “Direction selectivity in the retina: symmetry and asymmetry in structure and function,” Nat Rev Neurosci 13, 194–208 (2012).

41. K. P. Szatko, M. M. Korympidou, Y. Ran et al., “Neural circuits in the mouse retina support color vision in the upper visual field,” Nat Commun 11, 3481 (2020).

42. M. Ahmadlou, L. S. Zweifel, and J. A. Heimel, “Functional modulation of primary visual cortex by the superior colliculus in the mouse,” Nat Commun 9, 3895 (2018).

43. D. C. M. Henderson, J. R. Vianna, J. Gobran et al., “Longitudinal In Vivo Changes in Retinal Ganglion Cell Dendritic Morphology After Acute and Chronic Optic Nerve Injury,” Invest Ophthalmol Vis Sci 62, 5 (2021).

44. E. Akyol, A. M. Hagag, S. Sivaprasad et al., “Adaptive optics: principles and applications in ophthalmology,” Eye (Lond) 35, 244–264 (2021).

45. D. J. Wahl, P. Zhang, J. Mocci et al., “Adaptive optics in the mouse eye: wavefront sensing based vs. image-guided aberration correction,” Biomed Opt Express 10, 4757–4774 (2019).

46. Y. Jian, J. Xu, M. A. Gradowski et al., “Wavefront sensorless adaptive optics optical coherence tomography for in vivo retinal imaging in mice,” Biomed Opt Express 5, 547–559 (2014).

47. V. J. DePiero, Z. Deng, C. Chen, et al., “Transformation of Motion Pattern Selectivity from Retina to Superior Colliculus,” J Neurosci 44 (2024).

48. V. Porciatti, “Electrophysiological assessment of retinal ganglion cell function,” Exp Eye Res 141, 164–170 (2015).

49. L. R. Ferguson, J. M. Dominguez, 2nd, S. Balaiya, et al., “Retinal Thickness Normative Data in Wild-Type Mice Using Customized Miniature SD-OCT,” PLoS One 8, e67265 (2013).

50. T. H. Kim, R. Weimer, and J. Elstrott, “Cellular-resolution OCT reveals layer-specific retinal mosaics and ganglion cell degeneration in mouse retina in vivo,” Neurophotonics 13, 015006 (2026).

51. X. Xia, Y. Yao, J. Liang et al., “Evaluation of focus measures for the autofocus of line scan cameras,” Optik 127, 7762–7775 (2016).

52. S. S. Nikonov, R. Kholodenko, J. Lem et al., “Physiological features of the S- and M-cone photoreceptors of wild-type mice from single-cell recordings,” J Gen Physiol 127, 359–374 (2006).

