## Supplemental File for "Closed-loop optical optimization enables patterned retinal stimulation *in vivo* at cellular scales"

This file contains:

- Figures S1 to S11
- Tables S1 to S4
- Methods S1 to S3
- References

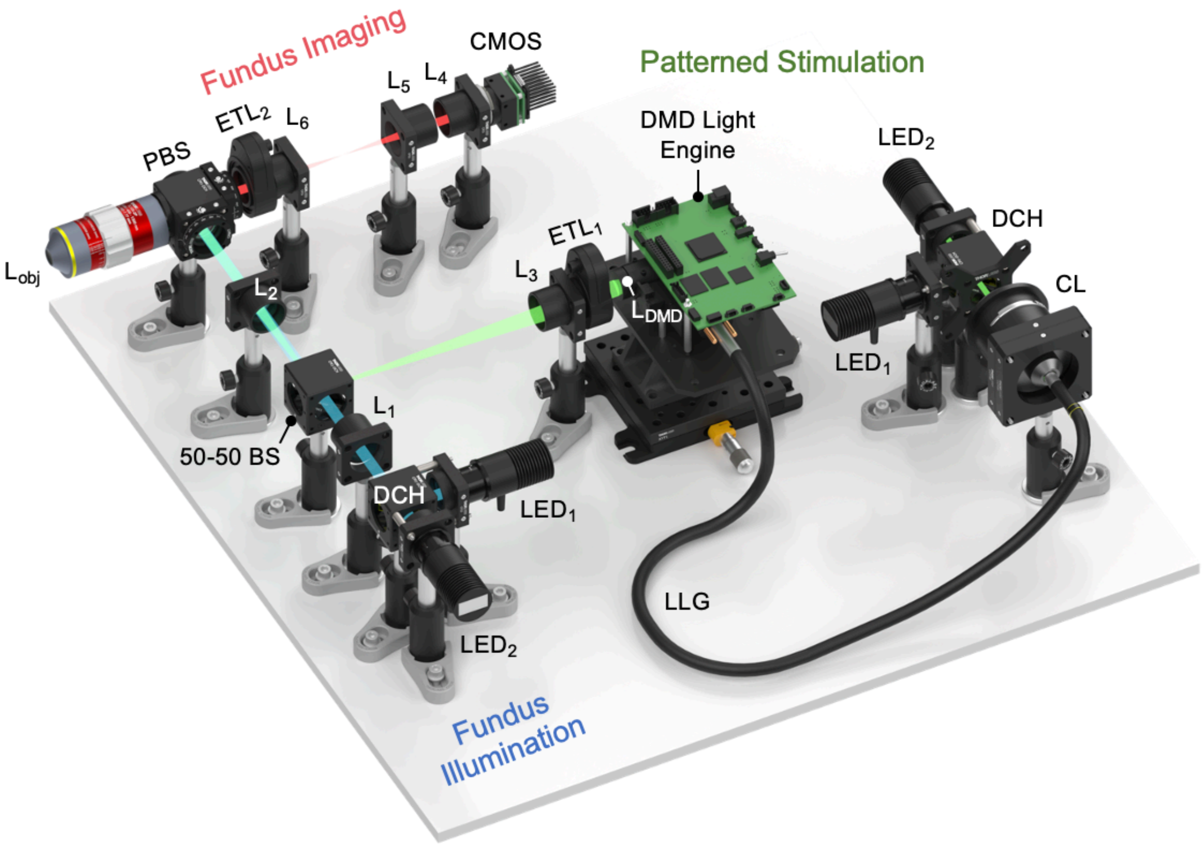


**Figure S1.** SolidWorks rendering of the experimental system. The layout illustrates the physical implementation of the fundus illumination, patterned stimulation, and fundus imaging pathways, corresponding to the schematic in Fig. 1(a).

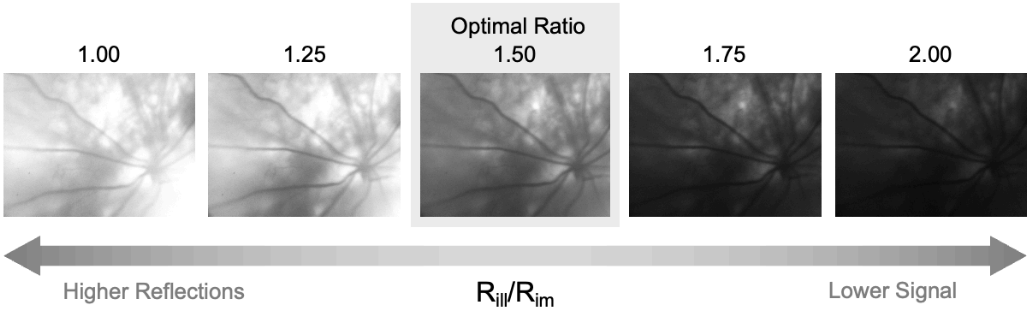


**Figure S2.** Effect of the pupil plane offset ratio $R_{ill}/R_{im}$ on fundus image quality. Increasing the ratio reduces corneal reflections, but also decreases the retinal signal. A value of $R_{ill}/R_{im} \approx 1.5$ provides the best tradeoff.

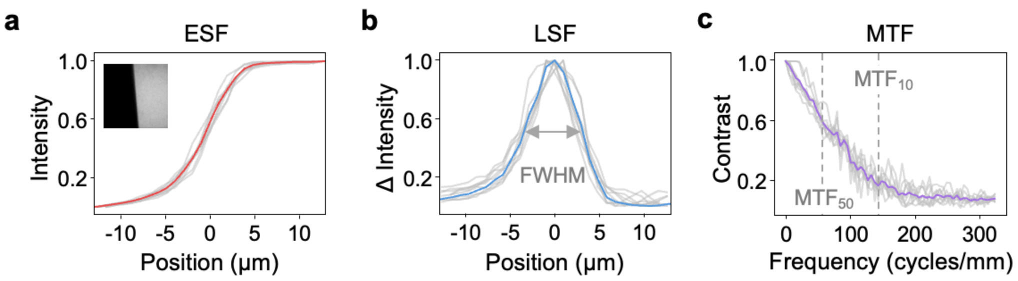


**Figure S3.** Quantification of $LED_2$ stimulus quality using the slanted-edge method. (a) Edge spread functions (ESFs) obtained by imaging a 10°-tilted edge projected from the digital micromirror device onto a camera positioned at the retinal conjugate plane using $LED_2$. The mean ESF is shown in red. (b) Corresponding line spread functions (LSFs) computed as the spatial derivatives of the ESFs, with the mean LSF shown in blue. The full width at half maximum (FWHM) of the LSF was taken as a direct measurement of the stimulus lateral resolution. (c) Modulation transfer functions (MTFs), obtained from the Fourier transform of the LSFs, with the mean MTF shown in purple. $MTF_{50}$ and $MTF_{10}$ denote the spatial frequencies at which contrast decreases to 50% and 10%, respectively.

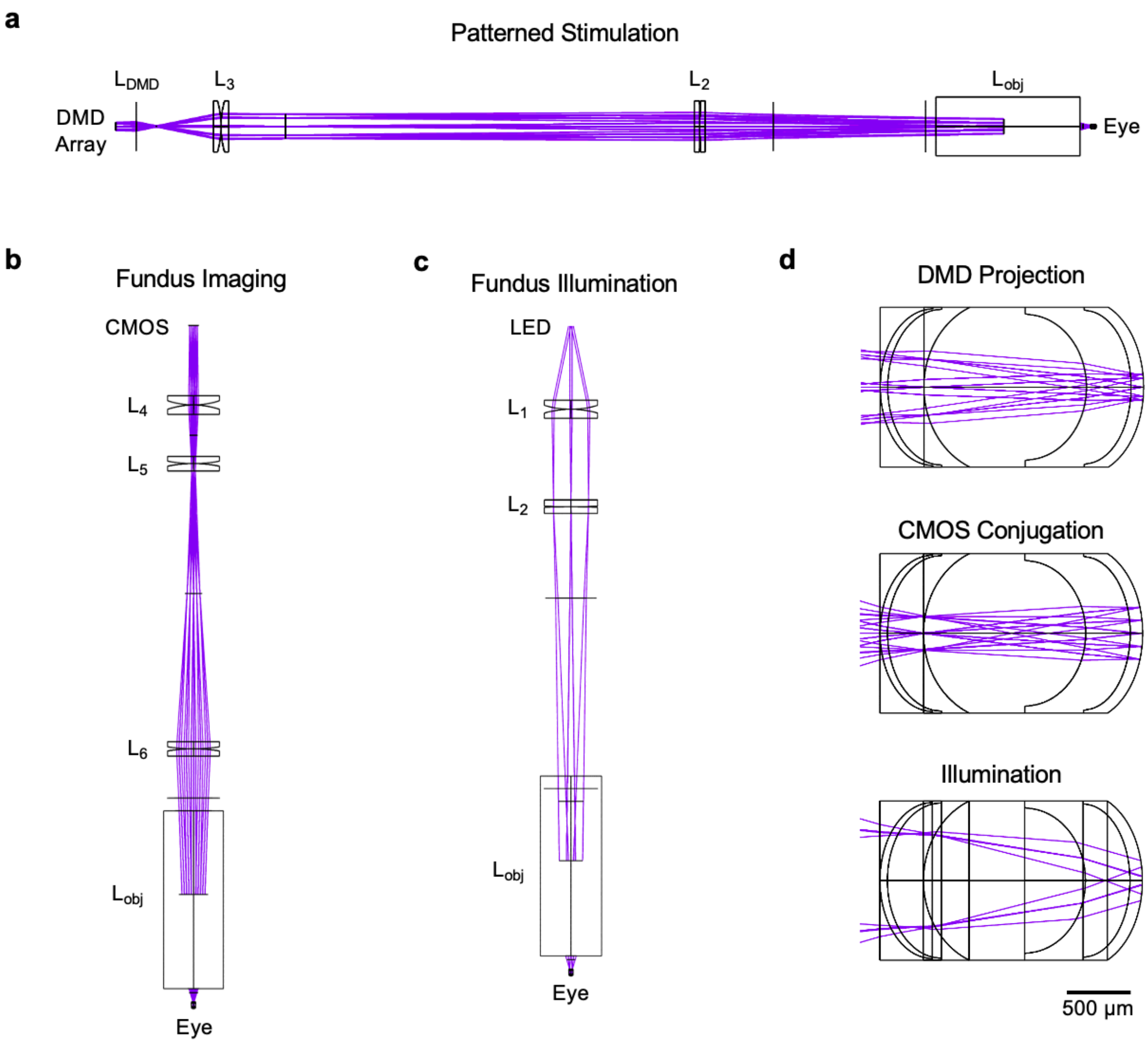


**Figure S4.** Sequential ray-tracing of the (a) patterned stimulation, (b) fundus imaging, and (c) fundus illumination pathways. (d) Magnified views illustrating the retinal conjugation of each pathway.

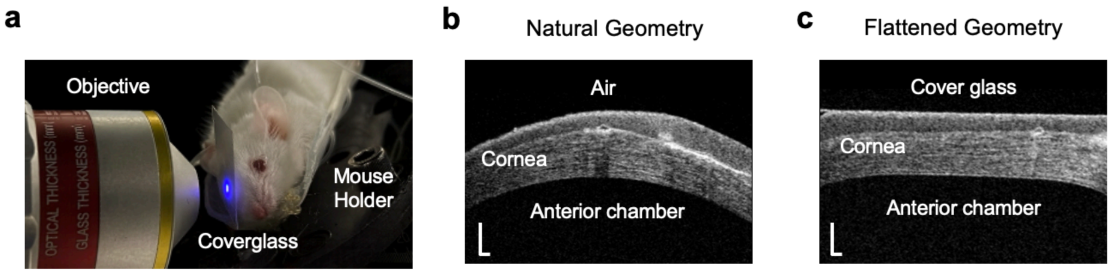


**Figure S5.** Effect of corneal flattening on ocular geometry and optical performance. (a) Photograph of the experimental setup showing application of a cover glass to the mouse cornea during stimulation. (b) Optical coherence tomography cross-sectional image of the cornea without cover glass. (c) Optical coherence tomography cross-sectional image of the cornea with cover glass. Scale bars: 50 µm.

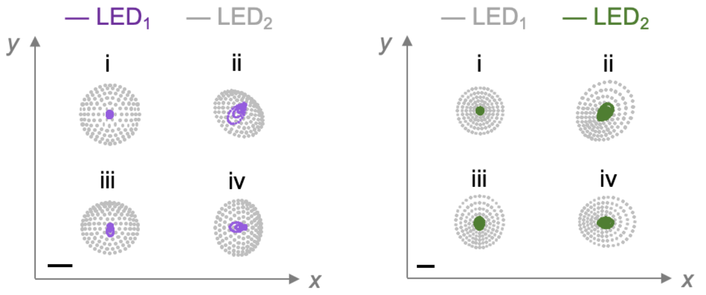


**Figure S6.** Ray-tracing results of a point stimulus delivered to the retina before application of the cover glass. The purple spots (left) represent point stimuli from $LED_1$ after proper mouse positioning, and the green spots (right) represent point stimuli from $LED_2$ after tunable lens correction. The gray spots correspond to point stimuli of the opposite LED under the same conditions. Spots were evaluated at the (i) center, (ii) corner, (iii) midpoint of the long edge, and (iv) midpoint of the short edge within a rectangular region of interest. Before the cover glass application, peripheral spots experienced major deviations due to coma and field curvature. Scale bars: 20 µm.

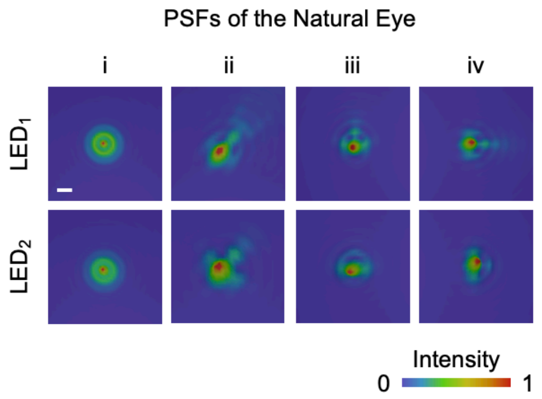


**Figure S7.** Point spread functions (PSFs) from four locations within the region of interest under the natural geometry of the mouse eye. Peripheral points (ii) - (iv) deviated from a Gaussian beam profile due to the aberrations of the eye.

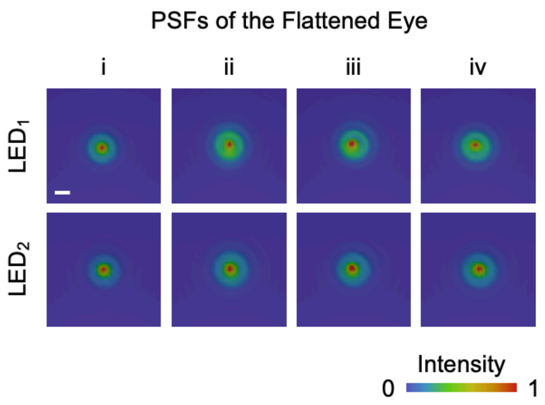


**Figure S8.** Point spread functions (PSFs) from four locations within the region of interest after application of the cover glass. Compared to those acquired with the natural optics of the mouse eye, the peripheral points (ii) - (iv) exhibited improved Gaussian characteristics.

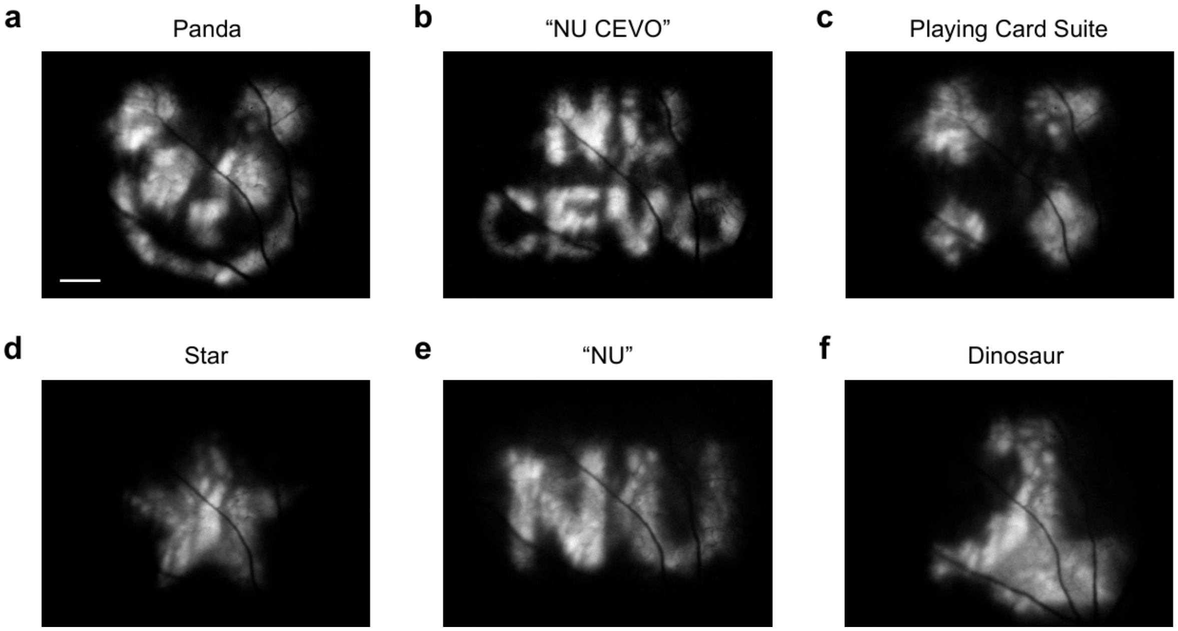


**Figure S9.** Representative patterned stimuli projected onto the retina at $\lambda_0$ = 415 nm. Examples include (a) a panda, (b) "NU CEVO" (Northwestern University Center for Engineering in Vision and Ophthalmology), (c) playing card suits, (d) a star, (e) "NU" (Northwestern University), and (f) a dinosaur. Scale bar: 120 μm.

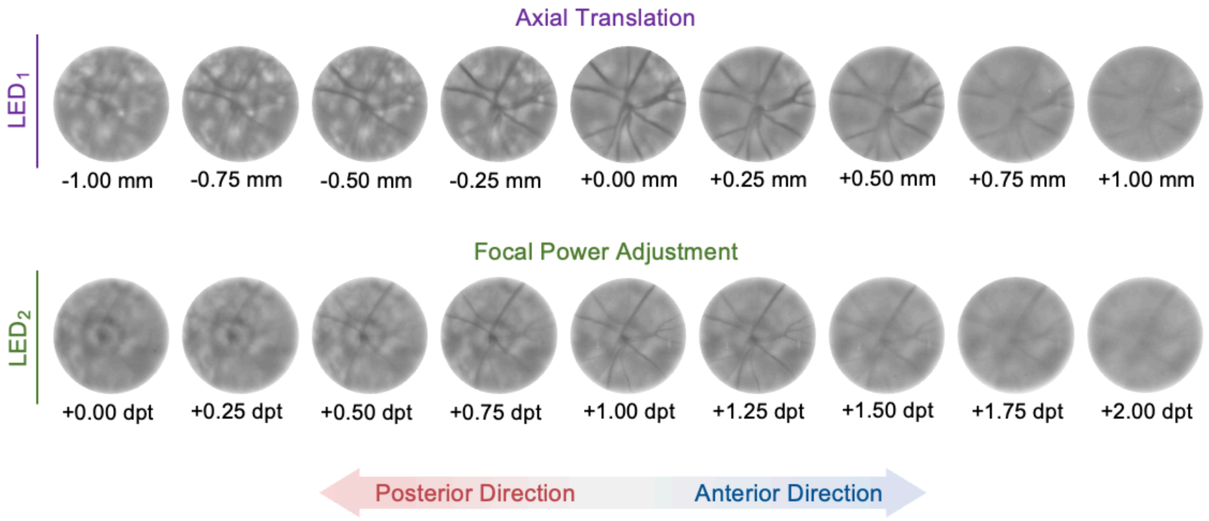


**Figure S10.** Retinal images acquired under different defocus conditions. The top row shows images obtained with $LED_1$ at varying axial positions, while the bottom row shows images acquired with $LED_2$ at varying dioptric offsets. Retinal vessels appear sharpest at the optimal focus and progressively blur with anterior and posterior defocus.

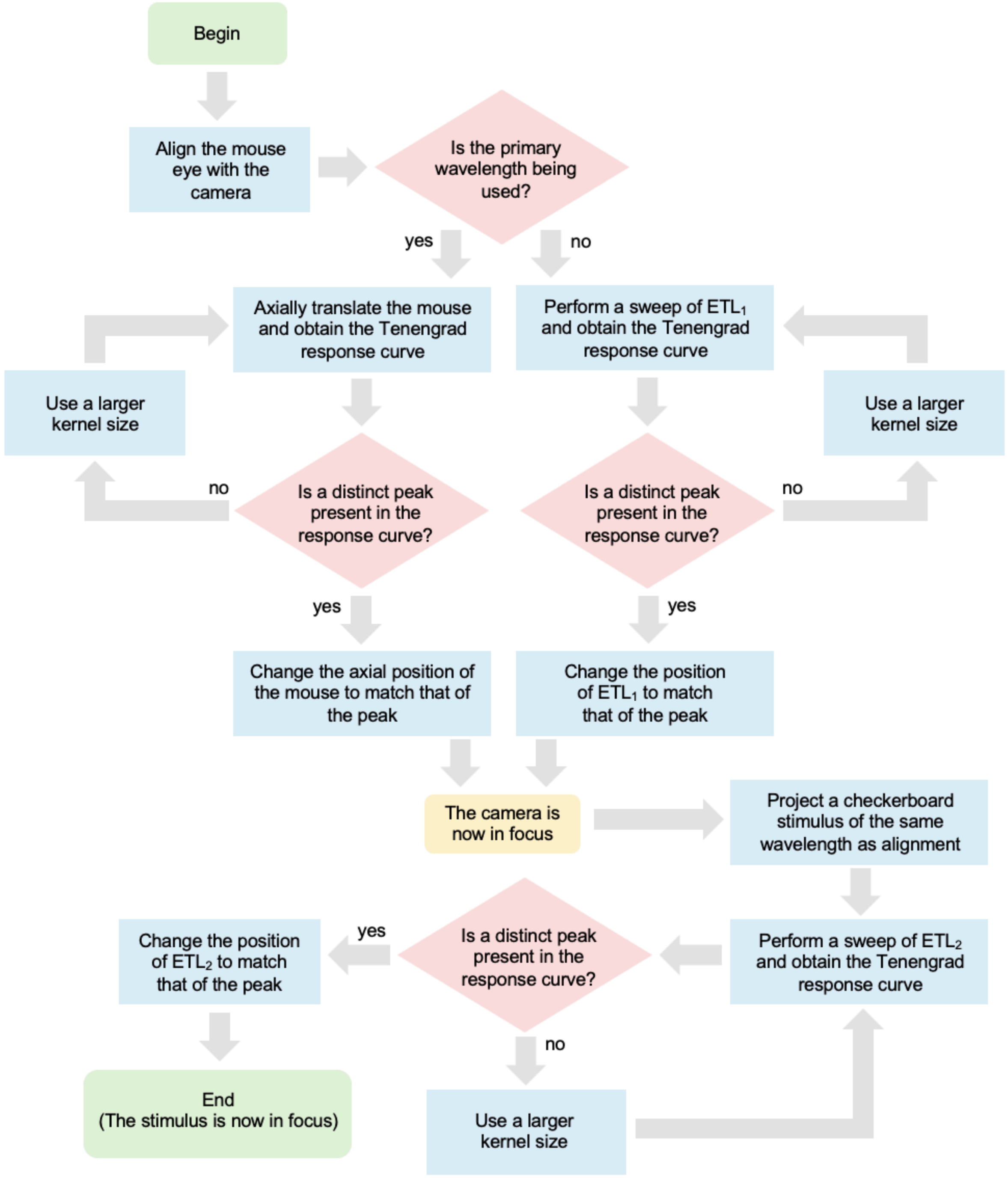


**Figure S11.** Flowchart of the closed-loop focus optimization workflow. Camera focus is initially established by manually translating the mouse using a translation stage, while chromatic focal shifts and stimulus focus are corrected using an automated ETL program.

**Table S1.** Measured lateral resolution and contrast metrics of the patterned stimulator.

| | $LED_1$ | $LED_2$ | Levene's Test p-value | t-Test | t-Test p-value |
|---|---|---|---|---|---|
| **FWHM** | 6.66 ± 0.45 µm | 6.44 ± 0.31 µm | 0.2287 | Student's | 0.2501 |
| **$MTF_{10}$** | 156.06 ± 11.27 cyc/mm | 153.87 ± 19.89 cyc/mm | 0.3405 | Student's | 0.7781 |
| **$MTF_{50}$** | 67.69 ± 5.80 cyc/mm | 67.89 ± 7.23 cyc/mm | 0.6125 | Student's | 0.9479 |

**Table S2.** Estimated values from checkerboard projection.

| Checker Size | $LED_1$ Relative Contrast | $LED_2$ Relative Contrast |
|---|---|---|
| **100 µm** | 0.93 | 0.97 |
| **50 µm** | 0.85 | 0.88 |
| **25 µm** | 0.83 | 0.84 |
| **12 µm** | 0.73 | 0.73 |
| **6 µm** | 0.50 | 0.51 |

**Table S3.** Ray-tracing spot diagram values (natural configuration).

| LED$_1$ Corrected | | | | |
|---|---|---|---|---|
| | **i** | **ii** | **iii** | **iv** |
| **RMS Radius at 415 nm** | 1.816 µm | 6.743 µm | 3.356 µm | 4.600 µm |
| **Geometric Radius at 415 nm** | 2.423 µm | 19.557 µm | 9.653 µm | 13.561 µm |
| **RMS Radius at 550 nm** | 19.602 µm | 15.891 µm | 17.938 µm | 17.125 µm |
| **Geometric Radius at 550 nm** | 25.614 µm | 23.491 µm | 24.895 µm | 24.345 µm |
| **LED$_2$ Corrected** | | | | |
| | **i** | **ii** | **iii** | **iv** |
| **RMS Radius at 415 nm** | 18.059 µm | 23.404 µm | 20.320 µm | 21.498 µm |
| **Geometric Radius at 415 nm** | 24.741 µm | 42.242 µm | 33.689 µm | 37.067 µm |
| **RMS Radius at 550 nm** | 1.816 µm | 6.167 µm | 3.718 µm | 4.669 µm |
| **Geometric Radius at 550 nm** | 3.102 µm | 15.209 µm | 9.164 µm | 11.554 µm |

**Table S4.** Ray-tracing spot diagram values (coverglass configuration).

| **$LED_1$ Corrected** | | | | |
|---|---|---|---|---|
| | **i** | **ii** | **iii** | **iv** |
| **RMS Radius at 415 nm** | 1.816 µm | 4.976 µm | 1.972 µm | 2.530 µm |
| **Geometric Radius at 415 nm** | 2.287 µm | 12.149 µm | 3.058 µm | 4.271 µm |
| **RMS Radius at 550 nm** | 16.452 µm | 18.425 µm | 16.706 µm | 17.226 µm |
| **Geometric Radius at 550 nm** | 19.763 µm | 25.820 µm | 25.472 µm | 26.113 µm |
| **$LED_2$ Corrected** | | | | |
| | **i** | **ii** | **iii** | **iv** |
| **RMS Radius at 415 nm** | 10.740 µm | 20.786 µm | 12.467 µm | 14.475 µm |
| **Geometric Radius at 415 nm** | 14.919 µm | 50.452 µm | 25.054 µm | 31.744 µm |
| **RMS Radius at 550 nm** | 1.816 µm | 4.784 µm | 3.070 µm | 3.638 µm |
| **Geometric Radius at 550 nm** | 2.153 µm | 9.340 µm | 6.288 µm | 7.872 µm |

**Methods S1.** Calculation of required power.

As described in the main text, the required radiant power to be delivered through the cornea is standardized for each center wavelength by the following equation.

$$P_c(\lambda_0) = \frac{E_{ret}A_{ret}}{T_{occ}(\lambda_0)S(\lambda_0)} \quad \text{(S1)}$$

where $E_{ret}$ denotes the retinal irradiance set to 1 μW/mm$^2$, $A_{ret}$ the retinal area being stimulated, $T_{occ}(\lambda_0)$ the ocular transmission, and $S(\lambda_0)$ the sensitivity at the specified wavelength. To calculate the stimulated area, we divide the dimensions of the DMD by the demagnification factor (4×) to obtain a field of 1.73 × 2.17 mm, corresponding to an area of $A_{ret}$ = 3.75 mm$^2$. However, if the field of view were to be cropped to 550 × 400 μm to achieve higher stimulus quality, as specified in the main text, the stimulated retinal area would change to $A_{ret}$ = 0.22 mm$^2$.

The ocular transmission $T_{occ}(\lambda_0)$ and cone sensitivities $S(\lambda_0)$ were estimated through literature [1, 2], a summary of which is shown in the plot below.

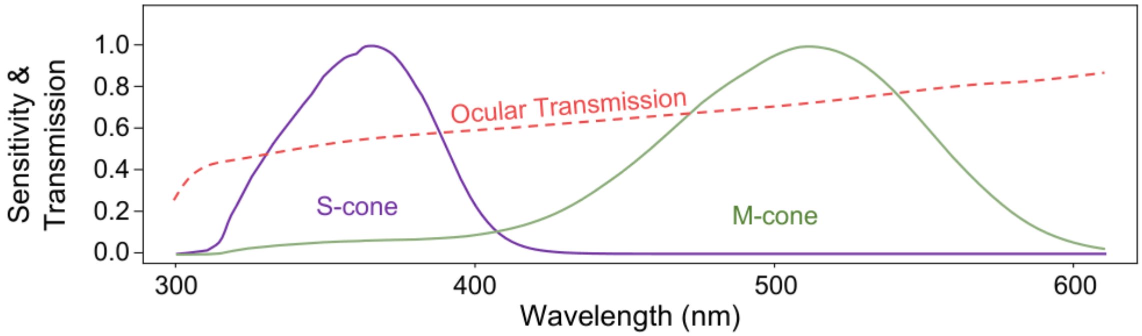


The corresponding transmission and cone sensitivity for $LED_1$ and $LED_2$ were found by evaluating these curves at their center wavelengths (415 nm and 550 nm, respectively), as summarized in the table below.

| Wavelength | $T_{occ}(\lambda_0)$ | $S(\lambda_0)$ |
|---|---|---|
| 415 nm | 0.60 | 0.11 |
| 550 nm | 0.82 | 0.40 |

Substituting these values into **Eq. (S1)**, we find that the output power measured immediately upon exiting the objective lens should be 56.8 μW for $LED_1$ and 11.4 μW for $LED_2$, assuming the entire stimulus field of view is used. If the stimulus were cropped to a 550 × 400 μm field of view, for instance, the power requirements would change to 3.33 μW for $LED_1$ and 0.67 μW for $LED_2$.

**Methods S2.** Calculation of average cone spacing.

We estimated cone spacing based on the literature-reported cone density of 12,400 cones/mm$^2$ [3]. The relationship between spacing $S$ and density $\rho$ can be approximated as

$$S \approx \sqrt{1/\rho}\,. \tag{S2}$$

Applying **Eq. (S2)** yields an estimated average cone spacing of 9.0 μm in the mouse retina. Because cone density decreases toward the retinal periphery and near the optic nerve head, the local cone spacing in these regions is expected to be larger.

**Methods S3**. Optimization of depth of focus.

Although a restrictive iris can, in principle, be placed at a pupil conjugate plane to match the stimulus depth of focus (DOF) to the axial length of the mouse eye, doing so comes at the expense of lateral resolution. We have designed the patterned stimulation pathway such that the DOF is large enough to span the distance between the superficial vascular plexus and the photoreceptor outer segments, yet small enough to maintain a lateral resolution comparable to the spacing between cone cells. Accordingly, lens $L_2$ has a diameter that serves as an aperture stop with NA = 0.05. We can approximate the DOF using

$$\mathrm{DOF} \approx \frac{\lambda}{\mathrm{NA}^2} + \frac{e}{\mathrm{M} \cdot \mathrm{NA}}, \qquad \text{(S3)}$$

where $\lambda$ is the wavelength, $e$ the pixel size, and M the magnification. The first term is the diffraction-limited axial resolution, while the second term describes the system limitations. Based on this calculation, stimuli from both LEDs experience a DOF greater than 180 μm, allowing the blood vessels of the superficial vascular plexus to serve as an indicator of the stimulus focal plane. At the same time, the lateral resolution is approximated using

$$\Delta x, y \approx \frac{0.61\lambda}{\mathrm{NA}}, \qquad \text{(S4)}$$

which remains below 7 μm for both LEDs. This allows vascular features to guide focus without sacrificing the stimulus's lateral resolution.

### References

1. Abballe L, Asari H. Natural image statistics for mouse vision. PLoS One. 2022;17(1):e0262763. https://doi.org/10.1371/journal.pone.0262763.
2. Henriksson JT, Bergmanson JP, Walsh JE. Ultraviolet radiation transmittance of the mouse eye and its individual media components. Exp Eye Res. 2010;90(3):382-7. https://doi.org/10.1016/j.exer.2009.11.004.
3. Jeon CJ, Strettoi E, Masland RH. The major cell populations of the mouse retina. J Neurosci. 1998;18(21):8936-46. https://doi.org/10.1523/JNEUROSCI.18-21-08936.1998.
